# Gray and White Matter Networks Predict Mindfulness and Mind Wandering Traits: A Data Fusion Machine Learning Approach

**DOI:** 10.1101/2024.04.15.589511

**Authors:** Minah Chang, Sara Sorella, Cristiano Crescentini, Alessandro Grecucci

## Abstract

**Objectives:** The main aim of our study was to find out which gray matter (GM) and white matter (WM) brain features are associated with mindfulness and mind wandering, and to investigate how mindfulness mediates deliberate and spontaneous mind wandering in terms of these associated brain components.

**Methods:** Structural MRI scans of 76 individuals and self-reported questionnaires were included in this analysis. We applied unsupervised machine learning algorithms to the brain imaging data to decompose the fused GM and WM into naturally grouping covarying networks. We then conducted a mediation analysis to assess the influence of mindfulness on deliberate and spontaneous mind wandering based on this data fusion networks. Additionally, we investigated if certain mindfulness facets mediate the two forms of mind wandering traits.

**Results:** We found GM and WM networks composed of structures that have been consistently linked to mindfulness (e.g., the cingulate, insula, basal ganglia) and fronto-parietal attentive regions exert direct effects on mindfulness, as well as on deliberate and spontaneous mind wandering. We also found an indirect mediating effect of the mindfulness facet *acting with awareness* on spontaneous mind wandering through one of the significant brain networks, we identified.

**Conclusions:** This study elicited the link between mind wandering and mindfulness, and expands our knowledge on the neural bases of these two psychological constructs.

## 1. Introduction

### 1.1 Background

Our attention is engaged in the present moment by immediate environmental stimuli that we perceive through the senses, and also by interoceptive cues such as hunger, pain, sleepiness, and sexual desire. On the other hand, our neurocognitive processes are not restricted to such immediacy (Mooneyham & Schooler, 2013; Smallwood & Schooler, 2015). Rather, the mind wandering away from conspicuous sensations and perceptions is considered the normal psychological baseline (Mason et al., 2007) during 25 to 50 percent of our waking hours (Kane et al., 2007; Killingsworth & Gilbert, 2010), especially when resting or sleeping, working, using a home computer, and commuting or traveling (Killingsworth & Gilbert, 2010). The self-generated thoughts of mind wandering that arise intrinsically within an individual can be either deliberate (intentional) or spontaneous (unintentional), and they can also be characterized as adaptive or maladaptive (Ottaviani & Couyoumdjian, 2013; Seli et al., 2016b).

Since the executive components of attention shift away when the mind wanders from the current environment, the mind wandering shares many similarities with traditional executive models of attention (Smallwood & Schooler, 2006; Smallwood & Schooler, 2015). According to control failure hypothesis, the mind wandering away from task-related thoughts could be a result of failure of executive control to keep on the current task (Kane & McVay, 2012; McVay & Kane, 2010). In support of this theory, the self-generation and disengagement of mind wandering are influenced by individual differences in executive control (Kane et al., 2007; Kane & McVay, 2012).

Mindfulness is largely comprised of two different components: self-regulation of attention, and adopting a particular orientation toward one’s experience in the present moment as characterized by curiosity, openness, and acceptance (Bishop et al., 2004; Messina et al., 2023; Monachesi et al., 2023). Therefore, mindfulness is also linked to attention and executive control as is mind wandering, though exerting its influence in the opposite direction. Structural evidence suggests that mindfulness training results in plasticity of brain mechanisms involved in emotional regulation through cognitive top-down emotion regulation (i.e., through the control of prefrontal cortex (PFC) to inhibit the influence of limbic areas, such as amygdala) in short-term training, but with perceptual bottom-up emotion regulation with long-term training (Brefczynski-Lewis et al., 2007; Chiesa et al., 2013; Taylor et al., 2011). The top-down emotion regulation in short-term training might be characterized as more deliberate cognitive control at a conscious level, and the bottom-up emotion regulation in long-term training might be characterized as more automatic control occurring below the level of conscious control. The latter would translate into a more stable trait that persists over time. Future studies may reveal the parallelism between top-down processes of mindful emotion regulation and deliberate mind wandering on the one hand, and bottom-up mindful emotion regulation and spontaneous mind wandering on the other.

Studies that examined the effects of short-term mindfulness training found that mind wandering tendencies can be significantly reduced even in as short as a two-week period (Feruglio et al., 2021; Mrazek et al., 2013a). Structural evidence shows that just 2 to 4 weeks of mindfulness training result in increased myelin and other axonal changes in white matters surrounding the anterior cingulate cortex (ACC) and posterior cingulate cortex (PCC) (Posner et al., 2014; Tang et al., 2010; Tang et al., 2012), and GM plasticity in the ventral PCC (Tang et al., 2020). In a meta-analysis study, Melis et al. (2022) reported that mindfulness training of over 6 weeks alters functional connectivity (FC) between networks involved in attention, executive function, emotional reactivity, and mind wandering. One study found that mindfulness training reduces default mode network (DMN) hyperconnectivity even after just one session (Zhang et al., 2023). These studies demonstrate plasticity of the brain that occurs in the early stages of mindfulness training, and raises the question of its link to the change in one’s mindfulness and mind wandering tendencies.

As for studies involving long-term mindfulness training, long-term meditators were compared to meditation-naïve controls. Across these studies, the ACC has been most consistently linked to mindful training, which is a region that has been found to be largely involved in mind wandering tendencies (Tang et al., 2015). Other regions consistently affected by mindfulness training across studies are known to be involved in meta-awareness (frontopolar cortex), body awareness (sensory cortices and insula), memory processes (hippocampus), self and emotion regulation (mid-cingulate cortex and orbitofrontal cortex along with ACC), and intra- and inter-hemispherical communication (superior longitudinal fasciculus and corpus callosum; Tang et al., 2015).

Studies have shown that there is a large number of parallels between DMN and mind wandering states, suggesting that they represent similar states of opposition to external perception and having anticorrelation to brain regions that engage in external sensory processes (Smallwood & Schooler, 2015). Meta-analysis of functional neuroimaging studies found that key DMN regions are also involved in mind wandering, including: medial PFC, PCC, medial temporal lobe, and bilateral inferior parietal lobule (Fox et al., 2015). Notably, a study by Mason et al. (2007) demonstrated that DMN recruitment is even greater during the periods with high-incidence of mind wandering. It has also been shown that medial PFC and PCC of DMN are relatively deactivated in experienced meditators compared to non-meditators. However, stronger connectivity has been found in mediators both at baseline and during meditation among the dorsolateral PFCs, dorsal ACC, and PCC, which are regions related to self-monitoring and cognitive control (Brewer et al., 2011).

Interestingly, the anterior PFC, the dorsal ACC, and the parahippocampal cortex were found to be more active during mind wandering when participants were unaware of their thoughts rather than aware (Christoff et al., 2009; Christoff et al., 2016). The evidence showing stronger recruitment of default and executive network regions in mind wandering outside of meta-awareness further suggests that spontaneous and deliberate mind wandering should be considered to be separate variables that engage different brain structures and functions. Previous findings also suggest that mindfulness traits should be treated as possible mediator in studies investigating mind wandering traits, and vice versa, as they share strong causal relations.

Most of the studies mentioned so far investigated mind wandering and mindfulness by comparing groups involved in different variations of mindfulness training that could have affected the subjects in different ways. The frequency of mind wandering in the lab and in daily life also have been found to share a positive correlation, indicating that the mind wandering tendency is a relatively stable individual trait (Hölzel et al., 2011; Murakami et al., 2012; Ottaviani & Couyoumdjian, 2013; Seli et al., 2016b; Taren et al., 2013; Zhuang et al., 2017). Indeed, it has been found that an eight-week mindfulness training induces significant positive change in three of the five individual traits associated with mindfulness defined in the Five Facets Mindfulness Questionnaire (FFMQ; Baer et al., 2006): *acting with awareness*, *nonjudging of inner experience,* and *observing* (Hozel et al., 2011). Mindfulness traits have been also associated with brain features. The FFMQ facet *describing* has been associated with the right anterior insula (Brodmann area, or BA 13), the right parahippocampal gyrus/amygdala (BA 28), the right dorsolateral PFC (BA 46), the right inferior parietal lobule (BA 40), and left superior PFC (BA 9) (Murakami et al., 2012; Zhuang et al., 2017); the FFMQ *nonjudging of inner experience* facet has been positively correlated with the surface area of the right superior PFC (BA 10); the *nonreactivity to inner experience* facet has been negatively correlated with the thickness of the right superior PFC (BA 8) and middle occipital cortex (Zhuang et al., 2017). In terms of brain function, mindfulness traits as measured by FFMQ were linked to increased FC among neural regions associated with attentional control, interoception, and executive function, and decreased FC among neural regions associated with self-referential processing and mind wandering (Parkinson et al., 2019). In addition, dispositional mindfulness of an individual has been found to be positively correlated with enhanced PFC and attenuated amygdala in emotion-regulation, which suggests that dorsomedial PFC regions are associated with mindfulness skills (Creswell et al., 2007; Modinos et al., 2010).

### 1.2 Current Study

The aim of this study was to model the associations among mindfulness, mind wandering, and morphometric features of the brain through a data fusion machine learning approach. In particular, we investigated whether or not deliberate and spontaneous mind wandering are mediated by one’s mindfulness characteristics. Many studies have so far put in much effort towards better understanding of different types of mind wandering and their neurological attributes. However, there has been no study up to date that has directed the investigation towards the mediating effect of mindfulness on mind wandering. The results of our study could also elucidate their possible inverse relationship, which has been a recurring question raised in previous studies. A clear implication from existing evidence is that mind wandering is influenced by DMN connectivity, and that the regions involved in these networks are susceptible to plastic change through mindfulness training that alters one’s mindfulness trait over time. In order to test the effect of mindfulness on mind wandering characteristics beyond existing inferences, we performed a mediation analysis of the trait variables and brain structure covariates composed of GM and WM networks.

In order to quantify and investigate the empirical relationship among mind wandering and mindfulness traits with associated brain regions, we obtained trait scores through respective self-report questionnaires along with structural brain data of our participants. For mind wandering, the scores for deliberate and spontaneous mind wandering were treated as separate variables as they engage different brain regions (Christoff et al., 2009; Christoff et al., 2016).

In recent years, we have witnessed the development and surge of machine learning, a computational intelligence model that builds algorithms to solve a specific task through exploration of patterns and reasoning. In cognitive science, machine learning algorithms are used in analyses such as neural decoding, neural response prediction, and hierarchical modeling (Fong et al., 2018). Computational approaches based on machine learning can quantify and decode brain network organization using pattern recognition, as well as perform predictive encoding of brain activities based on supervising metrics. Unlike previous structural studies that examined GM and WM alterations separately, we applied a data fusion approach to include both modalities in one model under the reasonable assumption that both GM and WM play important roles in mind wandering and mindfulness. The combinations of linked data points can capture the patterns of high-dimensional dataset, which univariate approach may fail to detect and miss significant findings (Calhoun & Sui, 2016). We applied unsupervised machine learning algorithms to the brain imaging data of our participants to decompose the fused GM and WM into naturally grouping covarying networks. We then conducted a mediation analysis to assess the influence of mindfulness on deliberate and spontaneous mind wandering based on this data fusion and network decomposition.

### 1.3 Aim and Hypothesis

The main aim of our study was to find out which GM and WM brain features are associated with mindfulness and mind wandering, and to investigate how mindfulness mediates deliberate and spontaneous mind wandering in terms of these associated brain components. To this end, after decomposing the brain into covarying GM and WM networks, we performed a mediation analysis to see if there are mediating effects of mindfulness on mind wandering traits. Additionally, we investigated if certain mindfulness facets mediate the two forms of mind wandering traits as an extra analysis for completeness. We predicted that specific GM and WM networks consisting of structures that have been consistently linked to mindfulness and mind wandering (such as the insula, the cingulate, the basal ganglia, and fronto-parietal attentive regions) would assert influence on mindfulness, which in turn would mediate deliberate and spontaneous mind wandering tendencies. We also predicted that some of the GM and WM networks would exert direct effects on mindfulness, and deliberate and spontaneous mind wandering.

## 2. Materials and Method

### 2.1 Participants

The data used in this study was entirely acquired from the MPI-Leipzig Mind-Brain-Body open-access database (Babayan et al., 2018), a project conducted by the Max Planck Institute of Human Cognitive and Brain Sciences in Leipzig, Germany, and approved by the ethics committee of the University of Leipzig. The project comprises of publicly available behavioral and brain imaging datasets of 318 participants with at least a structural quantitative T1-weighted image and a 15-minute resting-state fMRI data (for details of all available data, see Babayan et al., 2019 and also Mendes et al., 2019). All participants underwent medical screening to be eligible for MRI sessions, and also were screened for any past and present neuropsychological issues. From the database, we selected 76 participants (33 females, 43 males) for this study based on the inclusion criteria of age (between 20-45 years old, *m* = 27.1, *SD* = 5.08), right-handedness, negative drug test screening result, not having any history of neurological or psychiatric diagnosis, and availability of scores for the questionnaires (FFMQ and MW-D/MW-S scales). The entire study was conducted in German.

### 2.2 Mind Wandering and Mindfulness Questionnaires

#### Deliberate Mind Wandering (MW-D) and Spontaneous Mind Wandering (MW-S) Scales

Deliberate and spontaneous mind wandering scales are self-reported questionnaires that ask the subjects to assess their mind wandering tendencies in everyday life. A German adapted version of MW-D and MW-S scales (Carriere et al., 2013) were used to quantify the subjects’ deliberate and spontaneous mind wandering traits. MW-D and MW-S are separate and distinct traits that involve different degrees of self-control and mechanisms (Seli et al., 2014). For instance, MW-D is positively associated with *non-reactivity to inner experience* category of FFMQ, while MW-S is negatively associated with this facet (Seli et al., 2014). The original MW-D and MW-S scales are 4-item self-report questionnaires using Likert-scale of 1 to 7 (1: not at all true; 7: very true), and they were tested and validated in previous studies (Mrazek et al., 2013b; Seli et al., 2014; Seli et al., 2016a). The MW-D scale includes items that are related to intentional mind wandering, such as "I allow myself to get absorbed in pleasant fantasy". The MW-S scale includes items that are related to unintentional mind wandering, such as "I mind-wander even when I’m supposed to be doing something else" (Carriere et al., 2013). The scales in this study were adopted to a 5 Likert-scale (1: almost never; 5: very often).

#### Five Facet Mindfulness Questionnaire (FFMQ)

A German translated version of the original English FFMQ was used to measure the subjects’ mindfulness traits. The questionnaire had been developed based on previously existing mindfulness questionnaires, and empirically tested and validated (see Baer et al., 2006). The FFMQ is based on conceptualization of mindfulness as a multifaceted construct with five distinct facets represented by five categories: *acting with awareness* (8 items, 40 points), *describing* (8 items, 40 points), *nonjudging of inner experience* (8 items, 40 points), *nonreactivity to inner experience* (7 items, 35 points), and *observing* (8 items, 40 points), with a total of 39 items and 195 points using a Likert-scale of 1 to 5 (1: never or very rarely true; 5: very often or always true). The *acting with awareness* category of the FFMQ (*act_awareness*) measures one’s attentiveness traits with questions such as “When I do things, my mind wanders off and I’m easily distracted.” The *describing* category (*describe*) assesses one’s ability to describe and label one’s sensations, perceptions, thoughts, and feelings with statements such as “I am good at finding words to describe my feelings”. The *nonjudging of inner experience* category (*nonjudge*) measures the one’s tendency to judge one’s own inner experience with questions such as “I criticize myself for having irrational or inappropriate emotions”. The *nonreactivity to inner experience* category (*nonreact*) assesses the level of one’s reactive tendency with questions such as “I perceive my feelings and emotions without having to react to them”. Lastly, the *observing* category of the FFMQ (*observe*) measures one’s self-reported propensity to observe, notice, and attend to sensations, perceptions, thoughts, and feelings by answering statements such as “When I am walking, I deliberately notice the sensations of my body moving”.

#### Behavioral Data Analysis

The behavioral data was analyzed with JASP (JASP Team, 2023), a platform based on R (R Core Team, 2021) programming language for statistical computing and graphics. To find out if there are significant differences based on demographic characteristics, we compared the participant groups based on gender and age groups for each of the mind wandering (MW-D and MW-S) and mindfulness facet (*act_awareness*, *describe*, *nonjudge*, *nonreact*, and *observe*) scores. Pearson’s correlation coefficients were then calculated to evaluate the relationship among the seven variables.

### 2.3 MRI Data Acquisition/Pre-processing

The T1-weighted images were acquired on a 3T Siemens Magnetom Verio Scanner equipped with a 32-channel head coil using a MP2RAGE sequence (TR = 5,000 ms, TE = 2.92 ms, TI1 = 700 ms, TI2 = 2,500 ms, flip angle 1 = 4°, flip angle 2 = 5°, voxel size = 1.0 mm isotropic, duration = 8.22 min). The images were pre-processed with SPM12 (Statistical Parametric Mapping; Penny et al., 2011) and CAT12 (Computational Anatomy Toolbox; Gaser et al., 2022) in the MATLAB (The MathWorks Inc., 2022) environment. A visual check of data quality was performed in order to identify any distortion, such as head motion and other artifacts. The images were oriented to anterior commissure as the origin, and segmented into gray matter (GM), white matter (WM), and cerebrospinal fluid. The processed images were registered with Diffeomorphic Anatomical Registration Through Exponentiated Lie algebra (DARTEL; Ashburner, 2007) toolbox for SPM12 and normalized to the MNI (Montreal Neurological Institute) space with a spatial Gaussian smoothing with 12-mm, full-width at half-maximum (FWHM) Gassian kernel.

### 2.4 Data Fusion and Network Decomposition Using Unsupervised Machine Learning

Pre-processed structural MRI data was decomposed into covarying networks using data-driven parallel independent component analysis (PICA; Liu & Calhoun, 2007), an unsupervised machine learning approach. PICA is a modified ICA that applies dynamic constraints to both GM and WM modalities simultaneously to assess their interrelationships, and decompose the brain into meaningful networks that share the same response pattern. The analysis takes into account both GM and WM structures (by fusing the two modalities) to determine canonical covariate patterns, which allows natural measure of dynamic brain functions and connectivity, and offers improved reliability compared to ICA estimates (with tighter clusters and differently bootstrapped data sets) with lower dimensionality. PICA was conducted with Fusion ICA Toolbox (FIT; http://trendscenter.org/software/fit) to fuse GM and WM to find covariate matrix, and decompose the brain into independent networks. This multi-model fusion technique identifies brain regions that covary in their properties under the reasonable assumption that they belong to networks involved in carrying out similar psychological functions. To assess the algorithmic reliability of estimated independent components, Icasso (see Himberg & Hyvärinen, 2003; Himberg et al., 2004) was run 10 times.

### 2.5 Mediation Analysis

We conducted mediation analyses to examine the causal relationship among mindfulness and mind wandering (MW-D and MW-S) facets with dependency on the PICA GM and WM networks. We assigned the GM-WM networks as the predictor variable, and tested the mind wandering and mindfulness facets as either mediator variable or outcome variable. The analysis was performed with JASP.

## 3. Results

### 3.1. Behavioral Data Analysis

For all the participants, the mean and standard deviation of the FFMQ scores were: *act_awareness* (*M* = 16.7 ± 2.82), *describe* (*M* = 28.8 ± 5.82), *nonjudge* (*M* = 20.8 ± 4.59), *nonreact* (*M* = 18.9 ± 3.54), and *observe* (*M* = 22.2 ± 3.41). For mind wandering scales, the scores were: MW-D (*M* = 3.5 ± 1.05), and MW-S (*M* = 3.2 ± 1.01). Correlations among the behavioral data obtained from the questionnaires were calculated with Spearman’s partial correlation. All correlations among the five variables from the FFMQ, MW-D, and MW-S are shown in Figure 1. Significant positive correlations were found between: *act_awareness* and nonjudge (*r_s_* = 0.300, *p* = .009), *act_awareness* and *observe* (*r_s_* = 0.348, *p* = .002), *observe* and MW-D (*r_s_* = 0.241, *p* = .039), and MW-D and MW-S (*r_s_* = 0.409, *p* < .001). Marginally significant positive correlation was also found between *act_awareness* and MW-S (*r_s_* = 0.221, *p* = .059). A significant negative correlation was found between *act_awareness* and MW-S (*r_s_* = -0.237, *p* = .019).

**Figure 1.**
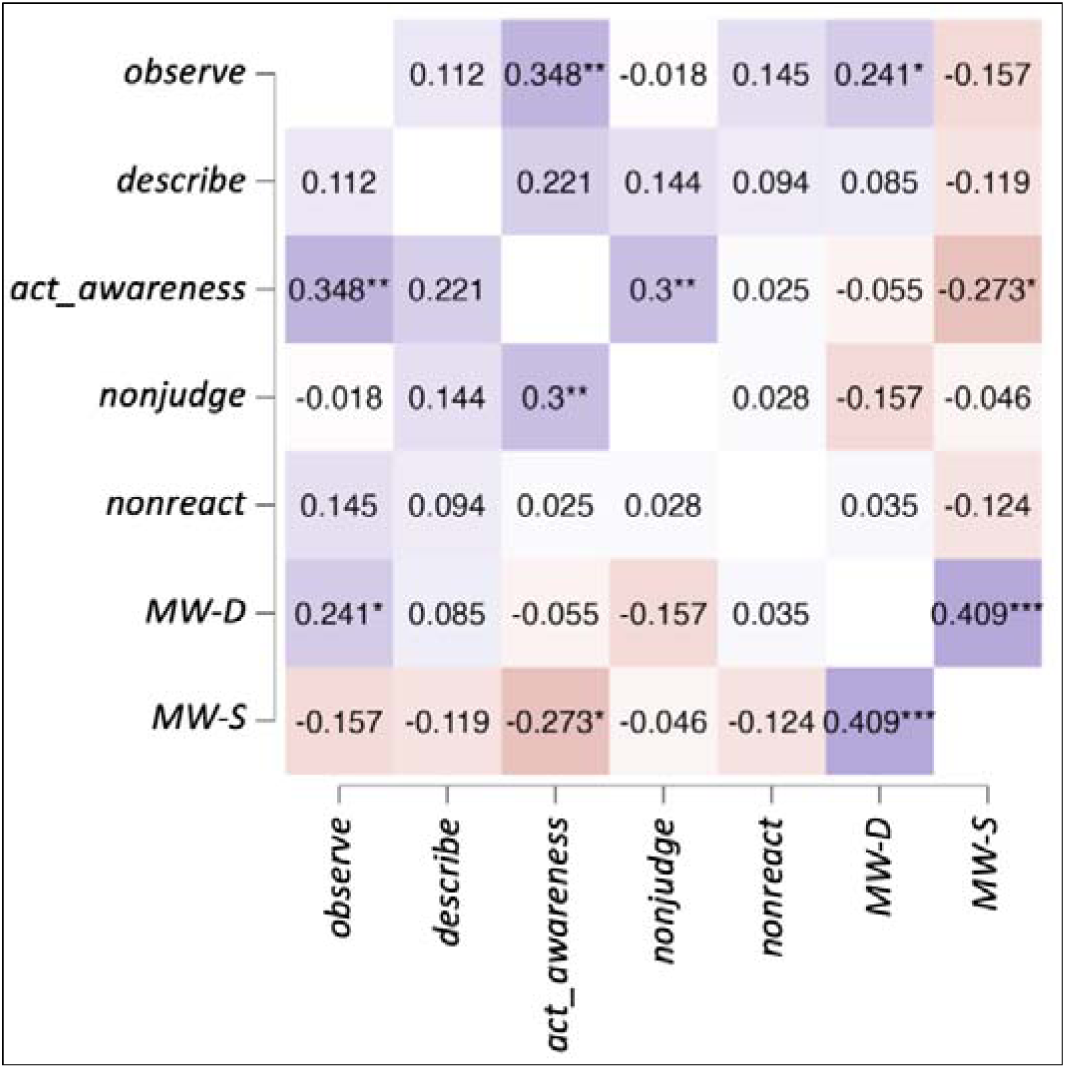
Spearman’s Partial Correlations Heatmap. **Fig. 1** shows all correlations among the five facets of FFMQ, MW-D, and MW. Significant values are denoted by * for *p* < .06, ** for *p* < .01, and *** for *p* < .001

We also preformed *t*-tests and correlations to evaluate the effects and gender and age for the FFMQ facets, MW-D, and MW-S (Table 1).

**Table 1.**
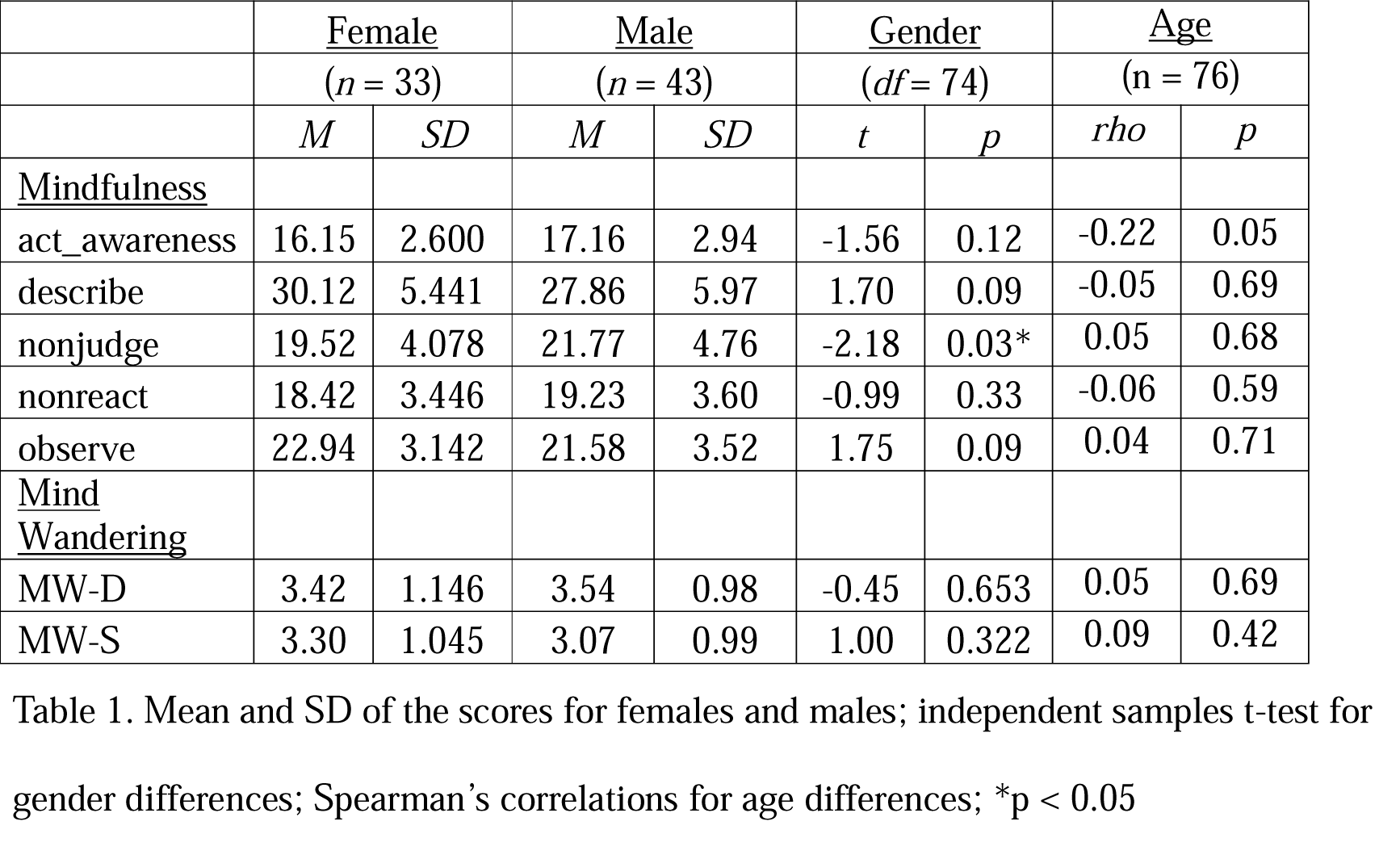
Effects of Age and Gender.

Mean and SD of the scores for females and males; independent samples t-test for gender differences; Spearman's correlations for age differences; \* $p < 0.05$
|  | <u>Female</u> |  | <u>Male</u> |  | <u>Gender</u> |  | <u>Age</u> |  |
| --- | --- | --- | --- | --- | --- | --- | --- | --- |
|  | (n = 33) |  | (n = 43) |  | (df = 74) |  | (n = 76) |  |
|  | *M* | *SD* | *M* | *SD* | *t* | *p* | *rho* | *p* |
| <u>Mindfulness</u> |  |  |  |  |  |  |  |  |
| act_awareness | 16.15 | 2.600 | 17.16 | 2.94 | -1.56 | 0.12 | -0.22 | 0.05 |
| describe | 30.12 | 5.441 | 27.86 | 5.97 | 1.70 | 0.09 | -0.05 | 0.69 |
| nonjudge | 19.52 | 4.078 | 21.77 | 4.76 | -2.18 | 0.03* | 0.05 | 0.68 |
| nonreact | 18.42 | 3.446 | 19.23 | 3.60 | -0.99 | 0.33 | -0.06 | 0.59 |
| observe | 22.94 | 3.142 | 21.58 | 3.52 | 1.75 | 0.09 | 0.04 | 0.71 |
| <u>Mind Wandering</u> |  |  |  |  |  |  |  |  |
| MW-D | 3.42 | 1.146 | 3.54 | 0.98 | -0.45 | 0.653 | 0.05 | 0.69 |
| MW-S | 3.30 | 1.045 | 3.07 | 0.99 | 1.00 | 0.322 | 0.09 | 0.42 |

There was an effect of gender for the mindfulness facet nonjudge (*t* = -2.18, *p* = 0.03). No other significant effect was found.

### 3.2 Data Fusion and Network Decomposition Using Unsupervised Machine Learning

The information theoretic criteria (see Wax & Kailath, 1985) estimated the presence of 18 independent covarying GM and WM networks in our data. The output from PICA consists of a matrix with the number of subjects (rows) and the loading coefficients for each PICA component (columns). The brain plots of all independent component estimates are shown in Figure 2 for the GM networks, and Figure 3 for the WM networks.

**Figure 2.**
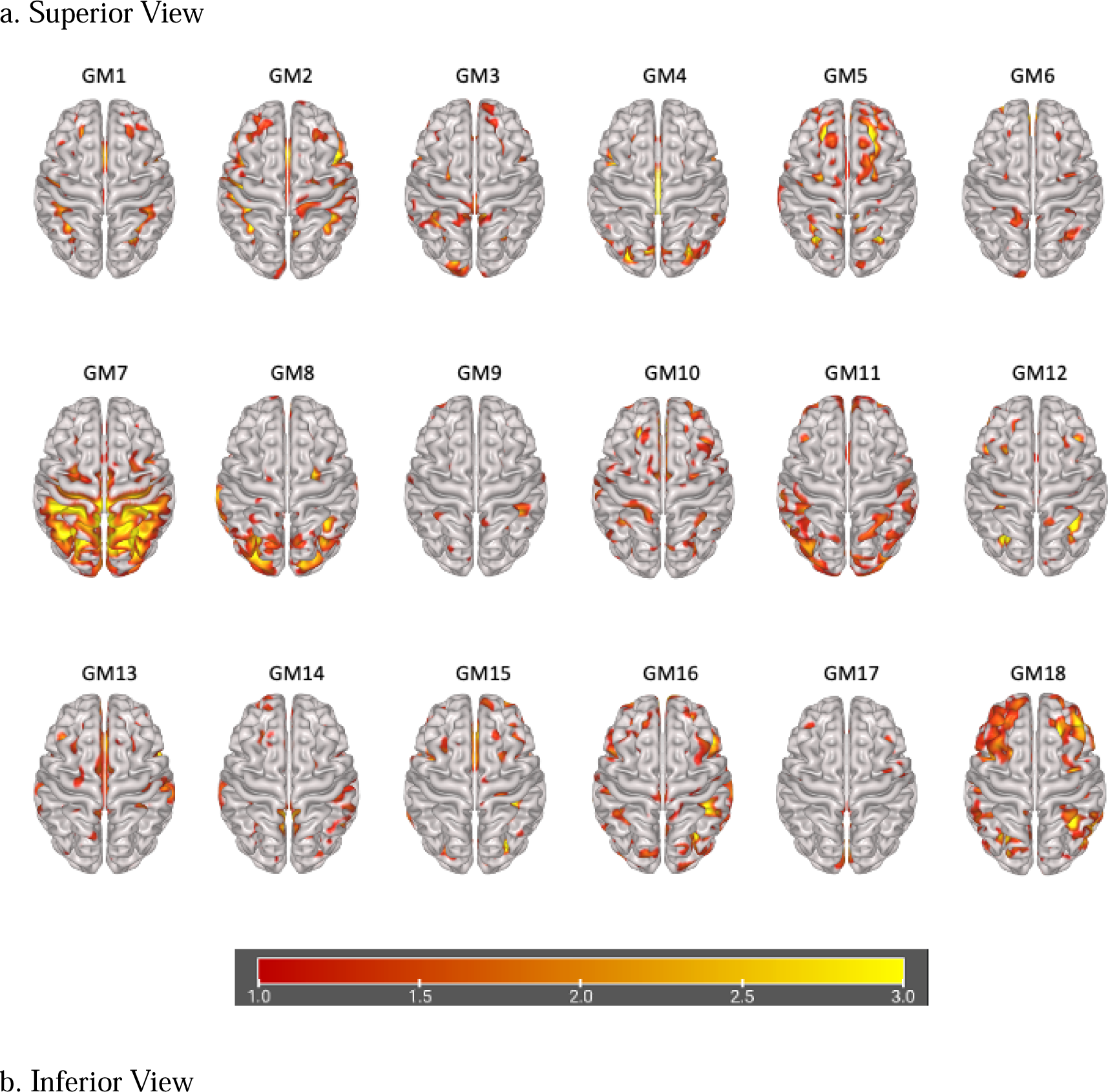
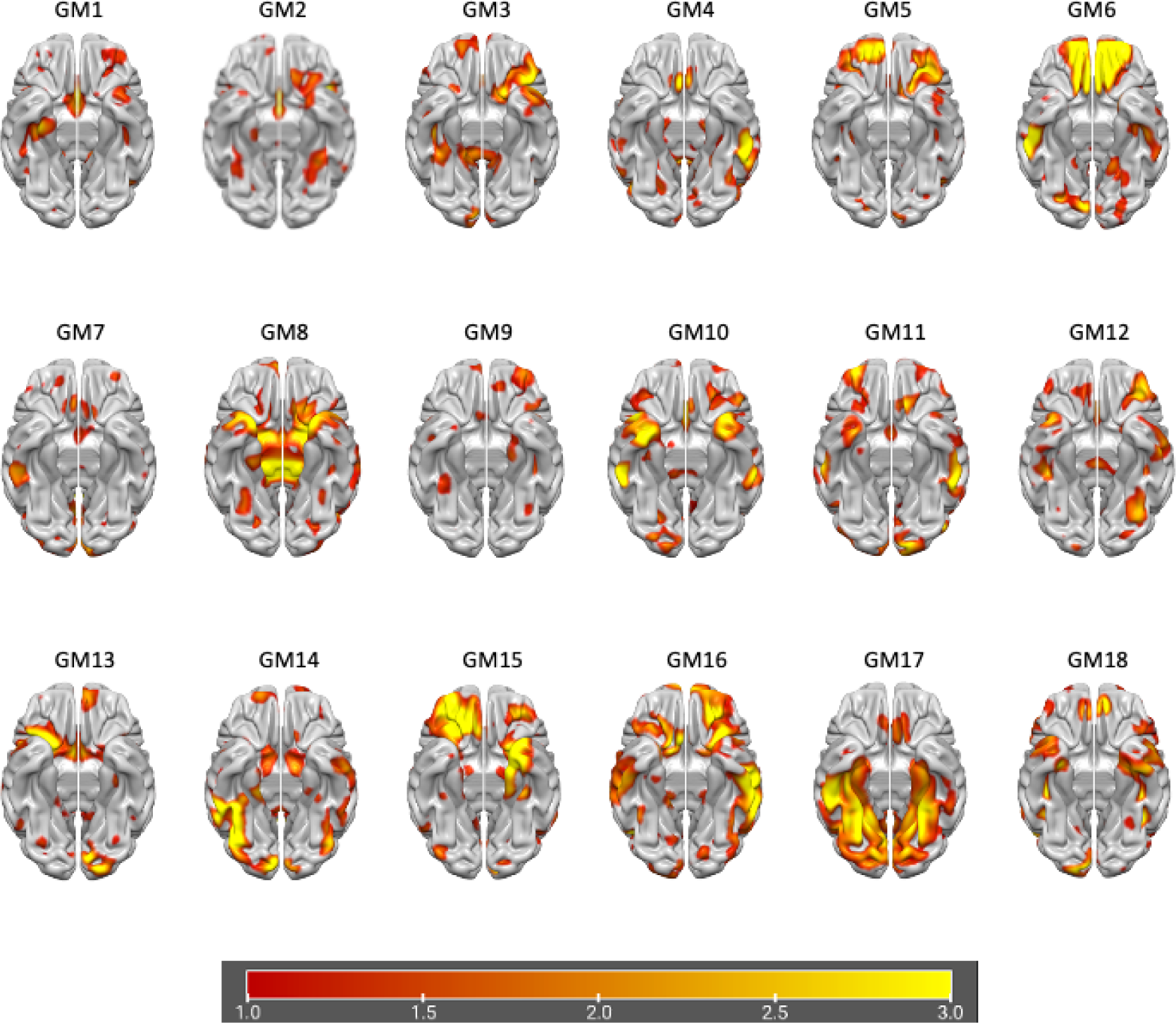
Plots of PICA GM Brain Networks. **Fig. 2** shows brain plots of the 18 GM independent covarying networks identified by unsupervised machine algorithms using the information theoretic criteria

**Figure 3.**
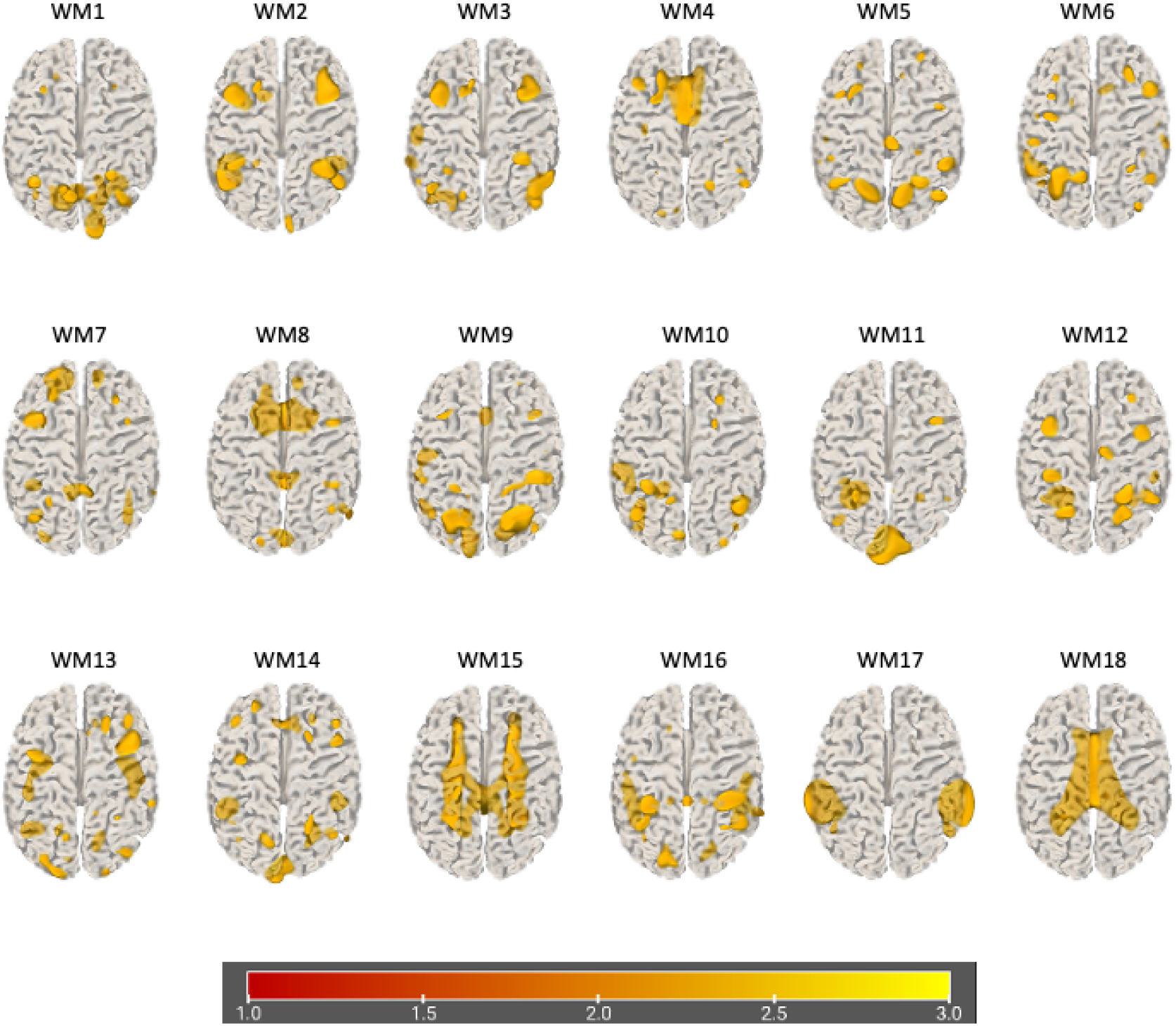
Plots of PICA WM Brain Networks (3d mesh) **Fig. 3** shows brain plots of the 18 WM independent covarying networks identified by unsupervised machine algorithms using the information theoretic criteria

### 3.3. Mediation Analysis

We conducted a mediation analysis to reveal which GM and WM networks are associated with the two forms of mind wandering (MW-D and MW-S), and if mindfulness plays a mediating role. In this analysis, the effect strength of the mediating variable (i.e., mindfulness measured with FFMQ scores) indicated the strength of its mediating influence among the predictor (i.e., GM and WM networks) and the outcome variables (i.e., MW-D and MW-S). Delta method approximation of standard errors was used to calculate the confidence interval with normal theory of confidence intervals and maximum likelihood estimator. Significant indirect effect indicates that there is an intervening variable that mediates the independent variable’s influence on the dependent variable (Biesanz et al., 2010). The summary of the significant findings reported in the following subsections are also summarized in Figure 4 with direct effects among the GM and WM networks, mindfulness, and mind wandering. Detailed morphometric features of the significant PICA networks are shown in the figures with lists of their positive and negative GM or WM concentrations in the corresponding tables beneath (Table 2, 3, 4, and 5). PICA components with a bilateral volume equal to or below 0.3 cc were removed from the tables in order to focus on more significant components that make up these networks.

**Figure 4.**
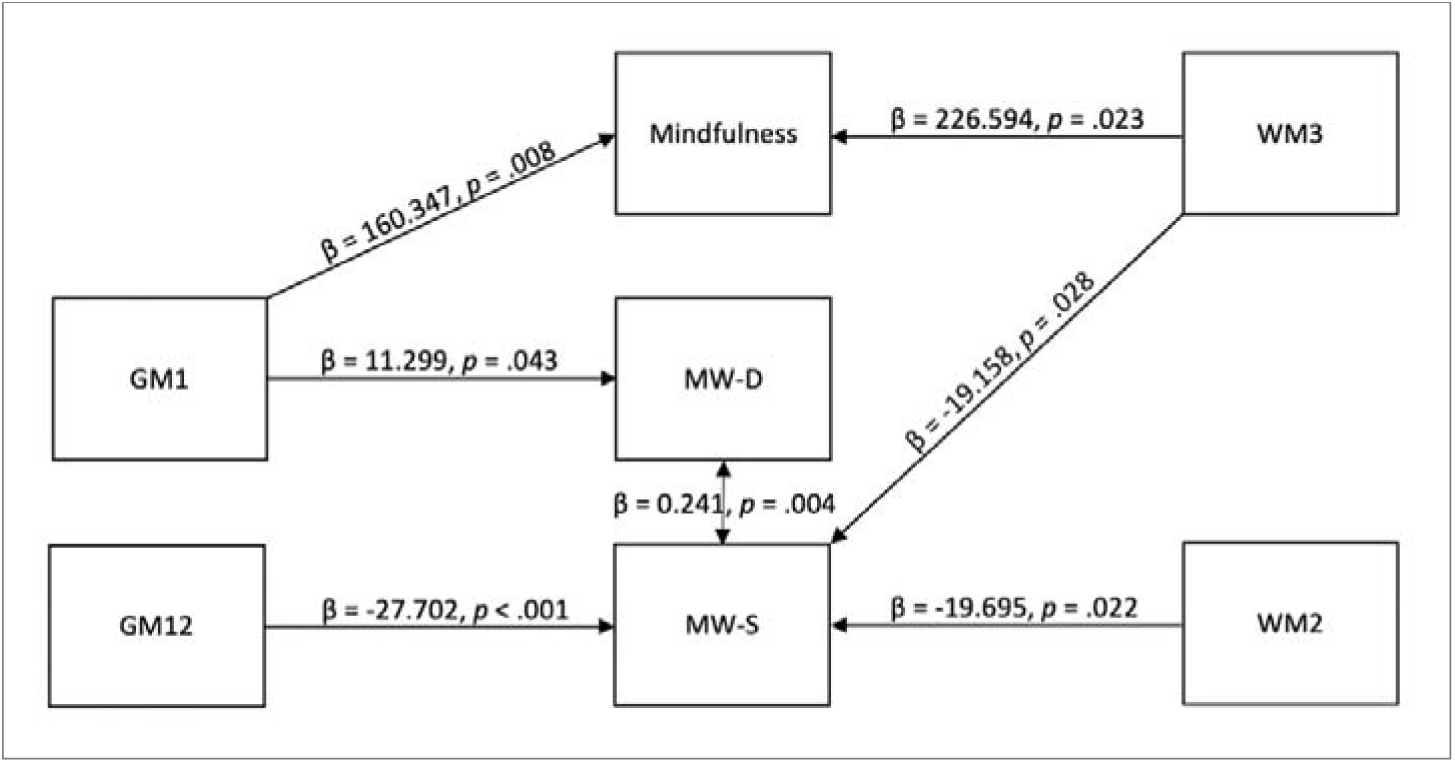
Direct Effects among GM and WM Networks, Mindfulness, and Mind Wandering. **Fig. 4** shows that GM1 has a direct effect on mindfulness and deliberate mind wandering, GM12 and WM2 have a direct effect on spontaneous mind wandering, WM3 has a direct effect on mindfulness and spontaneous mind wandering, and the two forms of mind wandering have a direct effect on each other

**Table 2.** GM1 Network Brain Components.

| <u>Area</u> | <u>Brodmann Area</u> | <u>Volume (cc)</u> | <u>Max value, MNI left/right (x, y, z)</u> |
| --- | --- | --- | --- |
| Caudate | 32 | 3.8/2.8 | 7.9 (-9, 12, 10)/6.4 (10, 8, 15) |
| Thalamus | 23, 30 | 1.5/1.5 | 6.9 (-10, -21, 16)/6.1 (13, -24, 15) |
| Caudate | 24, 32 | 4.4/3.7 | 6.7 (-3, 10, 5)/6.0 (12, 4, 18) |
| Thalamus | 30, 31 | 2.4/2.0 | 6.2 (-15, -28, 14)/5.6 (21, -35, 9) |
| Parahippocampal Gyrus | 30 | 0.3/0.1 | 5.6 (-24, -38, 5)/4.8 (24, -39, 3) |
| Hippocampus/Caudate | 30, 31 | 0.4/0.6 | 4.7 (-27, -38, 2)/4.4 (30, -36, 1) |
| Precuneus | 7 | 0.0/0.4 | 0 (0, 0, 0)/4.3 (18, -60, 43) |

*Note.* The first column indicates the name of the brain area, the second column indicates the name according to the Brodmann classification, the third column indicates the volume of gray or white matter concentration, and the fourth column indicates the coordinates of the peak for the area.

**Table 3.** GM12 Network Brain Components.

| Area | <u>Brodmann</u><br><u>Area</u> | <u>Volume</u><br><u>(cc)</u> | <u>Max value, MNI left/right (x, y, z)</u> |
| --- | --- | --- | --- |
| Cerebellar Tonsil | 37 | 0.5/3.3 | 4.9 (-37, -38, -38)/9.2 (49, -50, -40) |
| Broca/Visual Motor | 7, 45, 46 | 1.5/1.0 | 6.2 (-37, 20, 21)/8.0 (31, -52, 39) |
| Inferior Parietal Lobule | 7 | 0.0/0.7 | 0 (0, 0, 0)/6.2 (36, -53, 39) |
| Precuneus | 7 | 0.8/0.1 | 6.1 (-28, -62, 36)/4.2 (25, -50, 44) |
| Tuber | 19, 37 | 0.4/1.0 | 4.1 (-46, -74, -25)/5.9 (55, -47, -29) |
| Fusiform Gyrus | 18, 19 | 0.0/0.4 | 0 (0, 0, 0)/5.3 (27, -87, -18) |
| Inferior Semi-Lunar Lobule | 37 | 0.0/0.4 | 0 (0, 0, 0)/5.3 (53, -60, -35) |
| Culmen | 37 | 0.0/0.4 | 0 (0, 0, 0)/5.0 (50, -44, -29) |
| Declive | 19 | 0.1/0.6 | 3.8 (-46, -74, -22)/4.8 (31, -85, -18) |

*Note.* The first column indicates the name of the brain area, the second column indicates the name according to the Brodmann classification, the third column indicates the volume of gray or white matter concentration, and the fourth column indicates the coordinates of the peak for the area.

**Table 4.** WM2 Network Brain Components.

| <u>Area</u> | <u>Brodmann</u><br><u>Area</u> | <u>Volume</u><br><u>(cc)</u> | <u>Max value, MNI left/right (x, y, z)</u> |
| --- | --- | --- | --- |
| Middle Frontal Gyrus | 9, 46 | 1.7/2.0 | 9.1 (-37, 17, 28)/8.6 (37, 18, 31) |
| Superior Temporal Gyrus | 39 | 1.1/0.1 | 8.6 (-42, -51, 25)/3.6 (48, -55, 29) |
| Supramarginal Gyrus | 39, 40 | 1.5/0.8 | 8.0 (-45, -51, 27)/5.0 (48, -52, 32) |
| Supramarginal Gyrus | 7, 39 | 1.7/2.6 | 7.4 (-39, -48, 25)/7.8 (34, -40, 39) |
| Inferior Parietal Lobule | 7, 39 | 0.9/1.2 | 7.8 (-45, -48, 25)/7.4 (37, -42, 42) |
| Culmen | 20, 37 | 0.8/0.2 | 7.1 (-46, -39, -28)/4.4 (45, -41, -27) |
| Cerebellar Tonsil | 20, 37 | 0.6/0.8 | 6.9 (-43, -40, -35)/6.4 (43, -40, -37) |
| Precentral Gyrus | 9, 46 | 0.1/0.3 | 4.1 (-43, 19, 35)/6.3 (37, 21, 34) |

*Note.* The first column indicates the name of the brain area, the second column indicates the name according to the Brodmann classification, the third column indicates the volume of gray or white matter concentration, and the fourth column indicates the coordinates of the peak for the area. The software (FIT) computes WM nomenclature according to the adjacent GM components. Therefore, the areas reported for WM components should be referred to only as reference for the corresponding coordinates.

**Table 5.** WM3 Network Brain Components.

| <u>Area</u> | <u>Brodmann</u><br><u>Area</u> | <u>Volume</u><br><u>(cc)</u> | <u>Max value, MNI left/right (x, y, z)</u> |
| --- | --- | --- | --- |
| Middle Frontal Gyrus | 9, 46 | 1.9/1.5 | 9.5 (-37, 19, 32)/5.7 (40, 25, 25) |
| Precentral Gyrus | 9 | 0.6/0.1 | 8.1 (-37, 16, 35)/4.0 (36, 19, 34) |
| Visual<br>Association/Insula | 9, 19, 46 | 1.2/2.6 | 5.7 (-27, -70, -2)/6.3 (37, 22, 25) |
| Inferior Temporal<br>Gyrus | 19 | 0.2/0.4 | 4.8 (-42, -67, 2)/6.3 (43, -67, 1) |
| Superior Temporal<br>Gyrus | 37, 44 | 0.7/1.3 | 4.5 (-56, -13, 1)/6.3 (50, -50, 16) |
| Middle Occipital Gyrus | 19 | 0.2/0.4 | 4.6 (-37, -65, 3)/5.8 (40, -65, 3) |
| Inferior Parietal Lobule | 7 | 0.1/0.3 | 4.0 (-30, -42, 55)/5.1 (36, -34, 42) |
| Middle Temporal<br>Gyrus | 37, 39 | 0.1/0.3 | 3.5 (-55, -53, 11)/4.6 (43, -61, 27) |

*Note.* The first column indicates the name of the brain area, the second column indicates the name according to the Brodmann classification, the third column indicates the volume of gray or white matter concentration, and the fourth column indicates the coordinates of the peak for the area. The software (FIT) computes WM nomenclature according to the adjacent GM components. Therefore, the areas reported for WM components should be referred to only as reference for the corresponding coordinates.

#### Mediation Effect of Mindfulness on MW-D

No indirect effect of mindfulness on MW-D was found. However, significant direct effect parameter estimates were found for GM1 → MW-D (β = 11.299, *SE* = 5.576, *z* = 2.026, *p* = .043, 95% CI [0.37, 22.228]) and the path coefficient for GM1 → Mindfulness was significant (β = 160.347, *SE* = 60.405, *z* = 2.655, *p* = .008, 95% CI [41.955, 278.739]). The residual covariance for MW-D ↔ MW-S was also significant (β = 0.241, *SE* = 0.085, *z* = 2.844, *p* = .004, 95% CI [0.075, 0.407]).

#### Mediation Effect of Mindfulness on MW-S

No indirect effect of mindfulness (as measured by the sum of FFMQ scores) on MW-S was found. However, significant direct effect parameter estimates were found for GM12 → MW-S (β = -27.702, *SE* = 7.29, *z* = -3.8, *p* < .001, 95% CI [-41.99, -13.414]) and WM2 → MW-S (β = -19.695, *SE* = 8.611, *z* = -2.287, *p* = .022, 95% CI [-36.574, -2.817]). There was also a significant total effect for WM3 → MW-S (β = -19.158, *SE* = 8.716, *z* = -2.198, *p* = .028, 95% CI [-36.241, -2.075]) with significant path coefficient for WM3 → mindfulness (β = 226.594, *SE* = 99.469, *z* = 2.278, *p* = .023, 95% CI [31.638, 421.549]). The residual covariance for MW-D ↔ MW-S was also significant (β = 0.336, *SE* = 0.105, *z* = 3.2, *p* = .001, 95% CI [0.13, 0.542])

#### Mediation Effect of Individual Mindfulness Facets on Mind Wandering

For completeness of the analysis, we also ran an additional mediation analysis for each of the mindfulness facets for their influence on mind wandering. In this process, we found a significant indirect mediating effect for the FFMQ facet *acting with awareness*. The indirect effects involved the effect of GM1 on MW-S with *acting with awareness* trait as the moderator (GM1 → *acting with awareness* → MW-S; β = -6.050, *SE* = 2.538, *z* = -2.384, *p* = .017, 95% CI [-11.024, -1.076]). Significant path coefficients for this moderating relationship are illustrated in Figure 5 between GM1 and *acting with awareness* (β = 39.047, *SE* = 14.034, *z* = 2.782, *p* = .005, 95% CI [11.54, 66.553]), and between *acting with awareness* and MW-S (β = -0.155, *SE* = 0.034, *z* = -4.624, *p* < .001, 95% CI [-0.221, -0.089]). Meanwhile, the direct effect of GM1 → MW-S was not significant (β = 6.970, *SE* = 4.299, *z* = 1.621, *p* = .105, 95% CI [-1.456, 15.397]). See Figures 6, 7, 8, and 9 for the relative brain plots.

**Figure 5.**
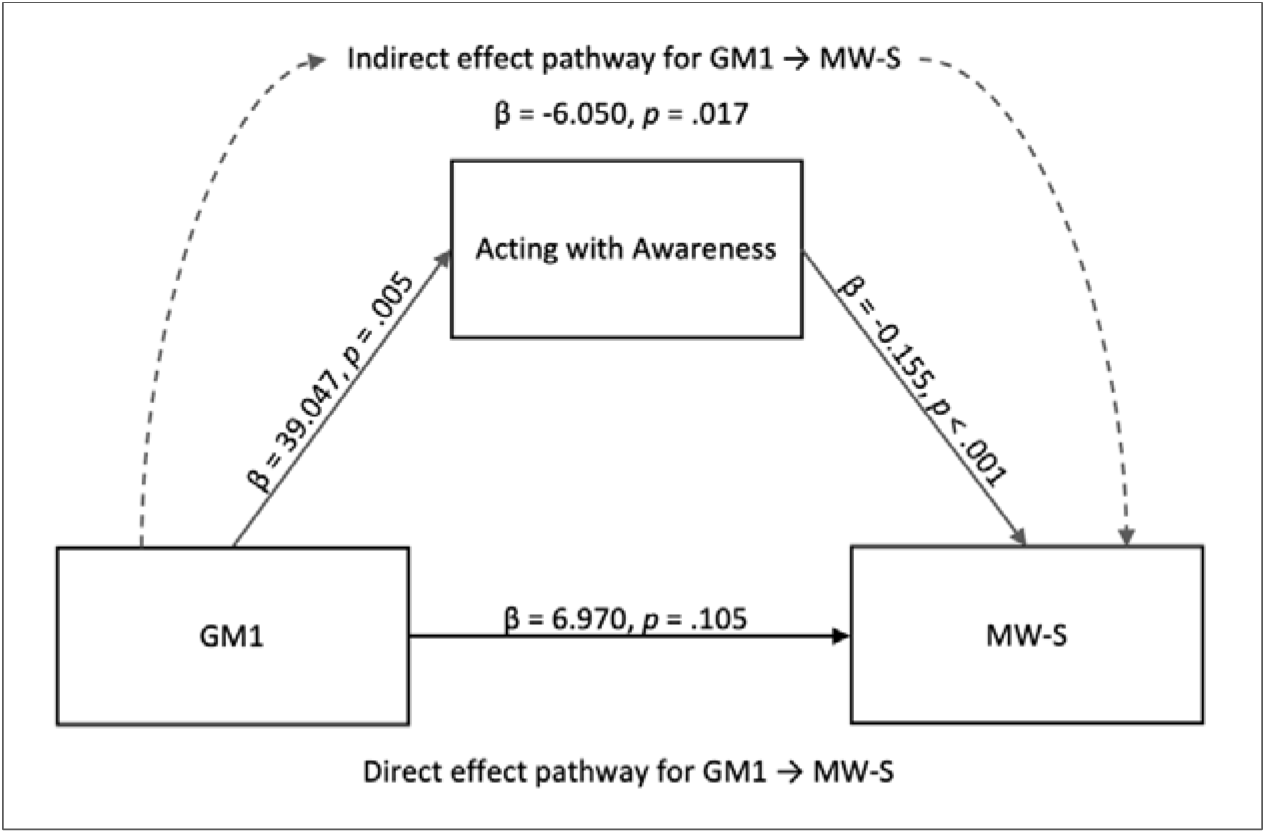
Mediation Model for the Indirect Effect of Acting with Awareness on MW-S. **Fig. 5** depicts the mediation effect of acting with awareness between GM1 component and spontaneous mind wandering

**Figure 6.**
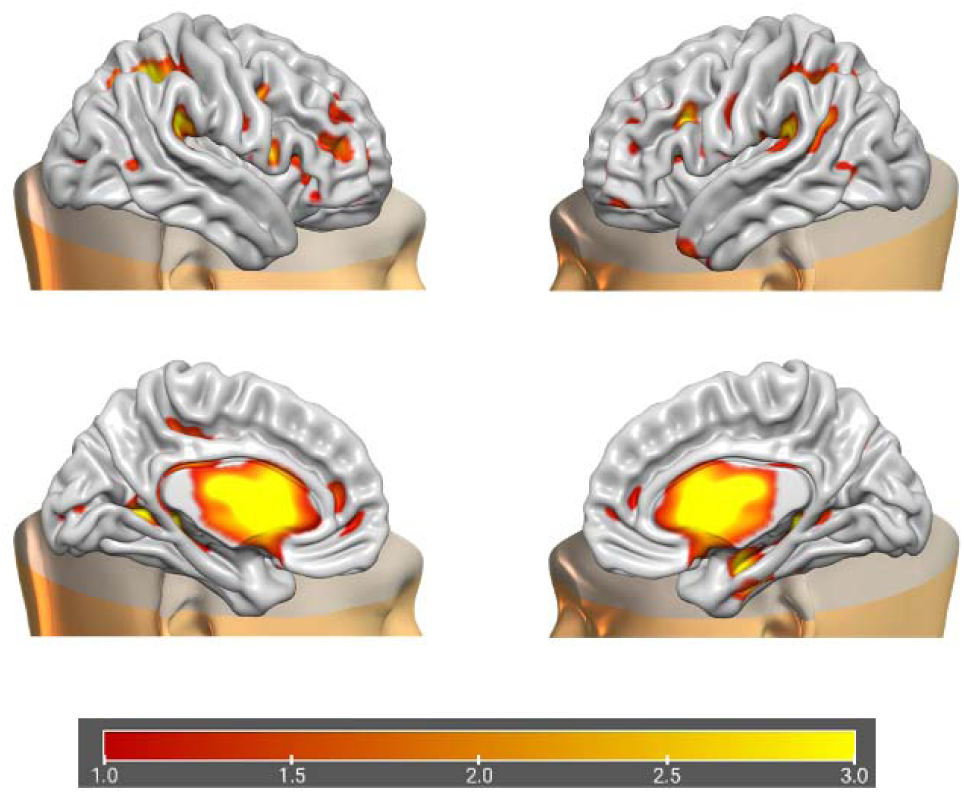
Morphometric Features of GM1. **Fig 6** shows the brain plots of GM1 components

**Figure 7.**
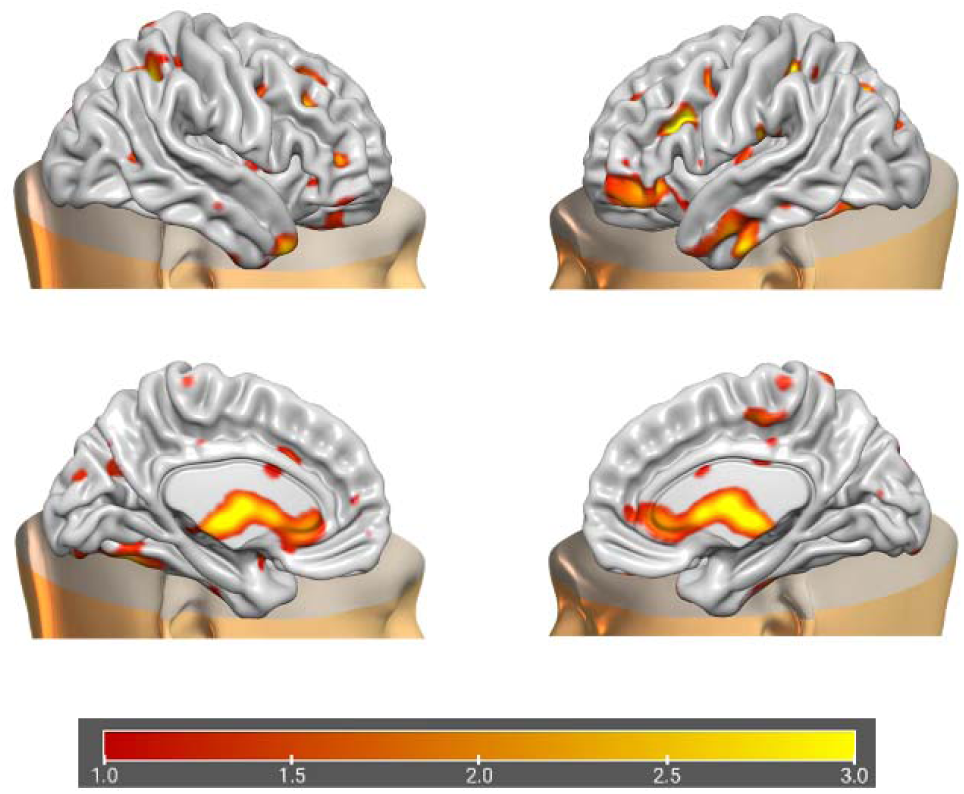
Morphometric Features of GM12. **Fig 7** shows the brain plots of GM12 components

**Figure 8.**
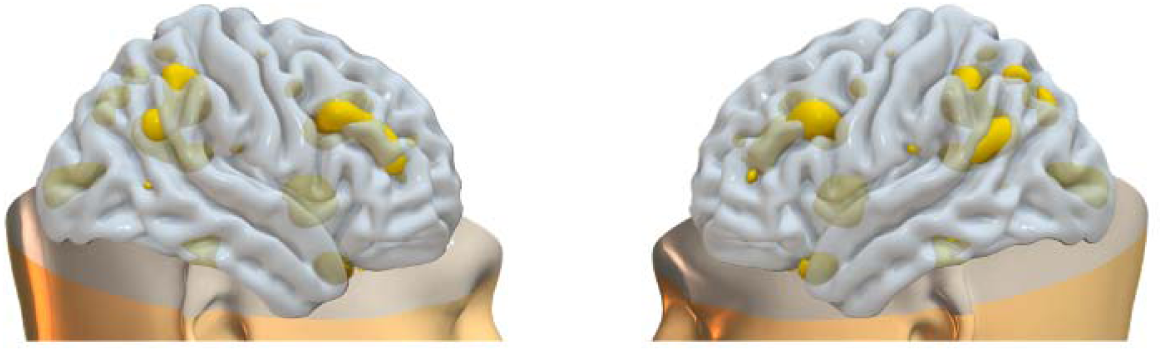
Morphometric Features of WM2. **Fig 8** shows the brain plots of WM2 components

**Figure 9.**
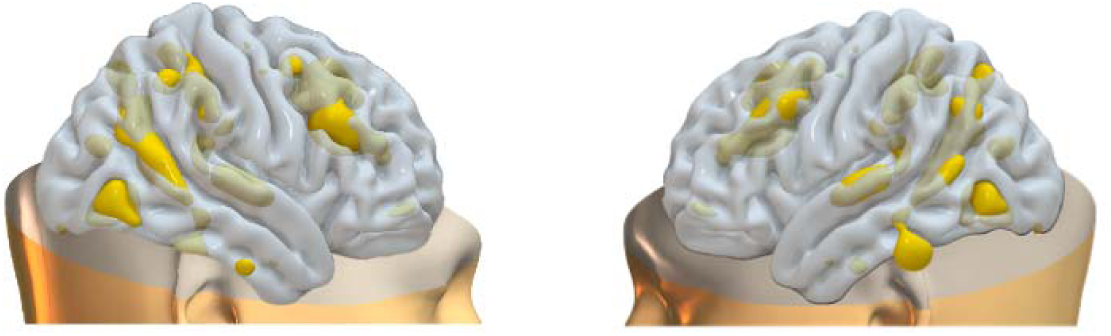
Morphometric Features of WM3. **Fig 9** shows the brain plots of WM3 components

## 4. Discussion

In the present study, our aim was to find morphometric brain features associated with mindfulness and mind wandering, and to investigate whether mindfulness mediates deliberate and spontaneous mind wandering in terms of these associated brain components. We found effective relationships among mindfulness, deliberate and spontaneous mind wandering, and PICA networks GM1, GM12, WM2, and WM3. In summary, PICA network GM1 was found to have a positive effect on deliberate mind wandering, GM12, WM2, and WM3 were found to have a negative effect on spontaneous mind wandering, and GM1 and WM3 were found to have a positive effect on mindfulness. In addition, WM1 had a positive relationship with the *acting with awareness* component of mindfulness, which was found to mediate spontaneous mind wandering. This mediation aligns with previous findings demonstrating a negative relationship between *acting with awareness* and spontaneous mind wandering (Sorella et al., 2025).

Our results further indicated a positive relationship between deliberate and spontaneous mind wandering. As a side note, GM1 correlated with WM5, and GM12 correlated with WM8 in the PICA fusion results; however, these WM counterparts were not significantly associated with the effects of interest in the mediation analysis. One possible explanation for this counterintuitive finding is that the correlations between GM1 and WM5, as well as GM12 and WM8, were negative. In the PICA fusion results, WM2 and WM3 were not correlated with any GM component.

A key finding of this study is the role of mindfulness – specifically the *acting with awareness* facet - in mediating the relationship between GM1 and spontaneous mind wandering. GM1 predominantly involves increased gray matter in subcortical structures, which supports previous research suggesting a top-down or bottom-up mindful emotion regulation mechanisms within limbic areas (Brefczynski-Lewis et al., 2007; Chiesa et al., 2013; Taylor et al., 2011). These findings also align with prior studies demonstrating gray matter changes in the hippocampus (Grant et al., 2010; Hölzel et al., 2008; Hölzel et al., 2011; Lu et al., 2014; Luders et al., 2009; Murakami et al., 2012), PCC (Hölzel et al., 2011; Lu et al., 2014), temporo-parietal junction (Hölzel et al., 2011), and cerebellum (Hölzel et al., 2011; Vestergaard-Poulsen et al., 2009) following mindfulness training. Previous studies also found that the caudate volume is negatively associated with dispositional mindfulness measured through the MAAS (Mindful Attention Awareness Scale; Brown and Ryan, 2003; Taren et al. 2013), and it was shown to be reduced through a 5-week mindfulness training in risky drivers (Mas-Cuesta et al., 2024). It has also been found that there is a negative relationship in terms of FC between the thalamus and PCC (measured through the MAAS scores), suggesting that these brain areas are involved in switching between mind wandering and mindfulness (Wang et al., 2014).

The GM1 network also includes reduced gray matter volume in the cingulate. However, we found a positive, rather than a negative relationship between the GM1 network and mindfulness scores. This could be explained by the fact that the mindfulness factor in our study is based on the FFMQ rather than the MAAS scores. Our results are also consistent with the findings of a meta-analysis study that show increased volume and activation of the thalamus, as well as increased brain activation of the caudate and in long-term meditators (Boccia et al. 2015). Results from other studies also support that mindfulness training results in increased volume of the caudate (Fahmy et al., 2018; Farb et al., 2012). The caudate is suggested to be involved in the alteration of habitual direction of attention (Baxter et al., 1992; Farb et al., 2012; McNab & Klingberg, 2008; Packard & Knowlton, 2002), while the thalamus is suggested to be involved in the orientation of attention and conscious awareness (Ward, 2013). These findings are coherent with our results showing positive relationship between GM1 and mindfulness. Furthermore, our mediation analysis shows that this network mainly involving the caudate and the thalamus is associated with *acting with awareness*, which can alter the process of mind wandering by influencing attentional processes. The GM12 network components also included brain structures previously discussed to be implicated in mindfulness and mind wandering, including the cerebellum, dorsolateral PFC, inferior parietal lobule, precuneus, and fusiform gyrus. The components are involved in mindfulness and mind wandering often through closely related functions such as self-referential processing, attentional processes, and other higher cognitive functions as well as interoceptive roles. Known modulatory role of the brain components in our significant networks well illustrate how these networks would directly or indirectly influence mindfulness and mind wandering facets in our study. **In terms of WM components,**

The maladaptive form of mind wandering can manifest as a cumbersome symptom in psychiatric diseases, and also exacerbated by factors such as aging, hyperactivity, time of the day, and depression (Giambra, 1995). Some studies even found negative correlation between mind wandering and happiness (Killingsworth & Gilbert, 2010; Grecucci et al., 2015; Mrazek et al., 2013b; Smallwood et al., 2004; Smallwood et al., 2007). Seneca expressed this notion in Letter to Lucilius (78, 14): “To be happy, two elements must be rooted out – the fear of future suffering, and the recollection of past suffering”. In this vein, studies suggest that the frequency of mind wandering in daily life might have an inverse relationship with mindfulness (Ottaviani & Couyoumdjian, 2013). Aforementioned studies have shown that mindfulness is positively linked to awareness (e.g., towards interoceptive and exteroceptive cues) and emotion regulation skills. We also mentioned that mindfulness training can improve a person’s mindfulness abilities after only a short session. Our study demonstrates that these changes can be represented by morphometric features of the brain to support behavioral reports, which can be useful in a clinical setting to measure short- and long-term results of a psychological intervention. As an example of practical application, a recent study showed that reduced activation of left cuneus is linked to positive outcome of cognitive behavior therapy across many studies, along with decreased activity in medial PFC and ACC (Yuan et al., 2022). Recent study also found that temporal variability of the DMN can predict spontaneous mind wandering (Sorella et al., 2025), which could be utilized as a biomarker for interventions aimed at reducing spontaneous mind wandering through mindfulness training.

in a clinical setting if a target intervention is to lesson episodes of spontaneous mind wandering through intervention such as mindfulness practices.

## Conclusion

Our study provides novel insights into the morphometric brain features associated with mindfulness, deliberate mind wandering, and spontaneous mind wandering, as well as the mediating effects of the FFMQ facet *acting with awareness* on spontaneous mind wandering. The covarying GM and WM networks in this study consisted of many brain structures that were linked to mindfulness and mind wandering in previous studies, but it is still unclear how and which of these structural components exert influence on one’s mindfulness and mind wandering traits. Our study also showed for the first time how mindfulness affects spontaneous mind wandering in a general population sample, complementing whole-brain data-driven predictive modeling approaches that focus on group-level effects (Kucyi et al., 2023). Future studies should aim to investigate individual-level structural and functional brain changes to establish biomarkers for monitoring the effectiveness of psychological interventions. Additionally, longitudinal studies with experienced meditators and clinical populations would further elucidate the mediating effects of mindfulness on spontaneous mind wandering.

## Acknowledgments

This work was supported by the BIAL Foundation Grant for Scientific Research (No.244/22).

## List Abbreviations

ACC: anterior cingulate cortex
BA: Brodmann area
DMN: default mode network
FC: functional connectivity
FFMQ: five facet mindfulness questionnaire
GM: gray matter
ICA: independent component analysis
MAAS: Mindful Attention Awareness Scale
MRI: magnetic resonance imaging
PCC: posterior cingulate cortex
PFC: prefrontal cortex
MW-D: deliberate mind wandering
MW-S: spontaneous mind wandering
PICA: parallel independent component analysis
WM: white matter

## Declaration

### Ethics Approval

The project was approved by the ethics committee of the University of Leipzig. The procedures used in this study adhere to the tenets of the Declaration of Helsinki.

### Consent to Participate

Informed consent was obtained from all individual participants in the study.

### Data and/or Code Availability

The data used in this study was entirely acquired from the MPI-Leipzig Mind-Brain-Body open-access database (Babayan et al., 2018; https://openneuro.org/datasets/ds000221/versions/1.0.0).

### Authors’ Contribution Statements

All authors equally contributed to this paper.

## References

Ashburner, J. (2007). A fast diffeomorphic image registration algorithm. Neuroimage, 38(1), 95–113. 10.1016/j.neuroimage.2007.07.007.

Babayan, A., Baczkowski, B., Cozatl, R., Dreyer, M., Engen, H., Erbey, M., Falkiewicz, M., Farrugia, N., Gaebler, M., Golchert, J., Golz, L., Gorgolewski, K., Haueis, P., Huntenburg, J., Jost, R., Kramarenko, Y., Krause, S., Kumral, D., Lauckner, M., … Villringer, A. (2018). MPI-Leipzig Mind-Brain-Body. OpenNeuro [ds000221, version 00002]. 10.18112/openneuro.ds000221.v1.0.0.

Babayan, A., Erbey, M., Kumral, D., Reinelt, J. D., Reiter, A. M. F., Röbbig, J., Schaare, H. L., Uhlig, M., Anwander, A., Bazin, P. L., Horstmann, A., Lampe, L., Nikulin, V. V., Okon-Singer, H., Preusser, S., Pampel, A., Rohr, C. S., Sacher, J., Thöne-Otto, A., … Villringer, A. (2019). A mind-brain-body dataset of MRI, EEG, cognition, emotion, and peripheral physiology in young and old adults. Scientific Data, 6, 180308. 10.1038/sdata.2018.308.

Baer, R. A., Smith, G. T., Hopkins, J., Krietemeyer, J., & Toney, L. (2006). Using self-report assessment methods to explore facets of mindfulness. Assessment, 13(1), 27–45. 10.1177/1073191105283504.

Baxter, L. R., Schwartz, J. M., Bergman, K. S., Szuba, M. P., Guze, B. H., Mazziotta, J. C., Alazraki, A., Selin, C. E., Ferng, H.-K., Munford, P., & Phelps, M. E. (1992). Caudate glucose metabolic rate changes with both drug and behavior therapy for obsessive-compulsive disorder. Archives of General Psychiatry, 49(9), 681–689. 10.1001/archpsyc.1992.01820090009002

Biesanz, J. C., Falk, C. F., & Savalei, V. (2010). Assessing mediational models: Testing and interval estimation for indirect effects. Multivariate Behavioral Research, 45(4), 661– 701. 10.1080/00273171.2010.498292.

Bishop, S. R., Lau, M., Shapiro, S., Carlson, L., Anderson, N. D., Carmody, J., Segal, Z. V., Abbey, S., Speca, M., Velting, D., & Devins, G. (2004). Mindfulness: A proposed operational definition. Clinical Psychology: Science and Practice, 11(3), 230–241. 10.1093/clipsy.bph077.

Boccia, M., Piccardi, L., & Guariglia, P. (2015). The meditative mind: A comprehensive meta-analysis of MRI studies. BioMed Research International, 2015, 1–11. 10.1155/2015/419808.

Brefczynski-Lewis, J. A., Lutz, A., Schaefer, H. S., Levinson, D. B., & Davidson, R. J. (2007). Neural correlates of attentional expertise in long-term meditation practitioners. Proceedings of the National Academy of Sciences, 104(27), 11483– 11488. 10.1073/pnas.0606552104.

Brewer, J. A., Worhunsky, P. D., Gray, J. R., Tang, Y. Y., Weber, J., & Kober, H. (2011). Meditation experience is associated with differences in default mode network activity and connectivity. Proceedings of the National Academy of Sciences, 108(50), 20254– 20259. 10.1073/pnas.1112029108.

Brown, K. W., & Ryan, R. M. (2003). Mindful Attention Awareness Scale (MAAS) [Database record]. APA PsycTests. 10.1037/t04259-000.

Calhoun, V. D., & Sui, J. (2016). Multimodal fusion of brain imaging data: A key to finding the missing link(s) in complex mental illness. Biological Psychiatry: Cognitive Neuroscience and Neuroimaging, 1(3), 230–244. 10.1016/j.bpsc.2015.12.005.

Carriere, J. S. A., Seli, P., & Smilek, D. (2013). Wandering in both mind and body: Individual differences in mind wandering and inattention predict fidgeting. Revue Canadienne de Psychologie Expérimentale, 67(1), 19–31. 10.1037/a0031438.

Chiesa, A., Serretti, A., & Jakobsen, J. C. (2013). Mindfulness: Top-down or bottom-up emotion regulation strategy? Clinical Psychology Review, 33(1), 82–96. 10.1016/j.cpr.2012.10.006.

Christoff, K., Gordon, A. M., Smallwood, J., Smith, R., & Schooler, J. W. (2009). Experience sampling during fMRI reveals default network and executive system contributions to mind wandering. Proceedings of the National Academy of Sciences, 106(21), 8719– 8724. 10.1073/pnas.0900234106.

Christoff, K., Irving, Z. C., Fox, K. C., Spreng, R. N., & Andrews-Hanna, J. R. (2016). Mind-wandering as spontaneous thought: A dynamic framework. Nature Reviews Neuroscience, 17(11), 718–731. 10.1038/nrn.2016.113.

Creswell, J. D., Way, B. M., Eisenberger, N. I., & Lieberman, M. D. (2007). Neural correlates of dispositional mindfulness during affect labeling. Psychosomatic Medicine, 69(6), 560–565. 10.1097/PSY.0b013e3180f6171f.

Fahmy, R., Wasfi, M., Mamdouh, R., Moussa, K., Wahba, A., Wittemann, M., Hirjak, D., Kubera, K. M., Wolf, N. D., Sambataro, F., & Wolf, R. C. (2018). Mindfulness-based interventions modulate structural network strength in patients with opioid dependence. Addictive Behaviors, 82, 50–56. 10.1016/j.addbeh.2018.02.013.

Farb, N. A., Segal, Z. V., & Anderson, A. K. (2012). Mindfulness meditation training alters cortical representations of interoceptive attention. Social Cognitive and Affective Neuroscience, 8(1), 15–26. 10.1093/scan/nss066.

Feruglio, S., Matiz, A., Pagnoni, G., Fabbro, F., & Crescentini, C. (2021). The impact of mindfulness meditation on the wandering mind: A systematic review. Neuroscience and Biobehavioral Reviews, 131,313–330. 10.1016/j.neubiorev.2021.09.032.

Fong, R. C., Scheirer, W. J., & Cox, D. D. (2018). Using human brain activity to guide machine learning. Scientific Reports, 8(5397), 1–10. 10.1038/s41598-018-23618-6.

Fox, K. C. R., Spreng, R. N., Ellamil, M., Andrews-Hanna, J. R., & Christoff, K. (2015). The wandering brain: Meta-analysis of functional neuroimaging studies of mind-wandering and related spontaneous thought processes. NeuroImage, 111, 611–621. 10.1016/j.neuroimage.2015.02.039.

Gaser, C., & Dahnke, R., Thompson, P. M., Kurth, F., & Luders, E. (2022). CAT – A computational anatomy toolbox for the analysis of structural MRI data. BioRxiv. 10.1101/2022.06.11.495736.

Giambra, L. M. (1995). A laboratory method for investigating influences on switching attention to task-unrelated imagery and thought. Consciousness and Cognition: An International Journal, 4(1), 1–21. 10.1006/ccog.1995.1001.

Grant, J. A., Courtemanche, J., Duerden, E. G., Duncan, G. H., & Rainville, P. (2010). Cortical thickness and pain sensitivity in zen meditators. Emotion, 10(1), 43–53. 10.1037/a0018334.

Grecucci, A., Pappaianni, E., Siugzdaite, R., Theuninck, A., & Job, R. (2015). Mindful emotion regulation: Psychological and neural mechanisms. BioMed Research International, 2015, 670724. 10.1155/2015/670724.

Himberg, J., & Hyvärinen, A. (2003). Icasso: software for investigating the reliability of ICA estimates by clustering and visualization. In IEEE XIII Workshop on Neural Networks for Signal Processing (p. 259–268). 10.1109/nnsp.2003.1318025.

Himberg, J., Hyvärinen, A., & Esposito, F. (2004). Validating the independent components of neuroimaging time series via clustering and visualization. NeuroImage, 22(3), 1214– 1222. 10.1016/j.neuroimage.2004.03.027.

Hölzel, B. K., Carmody, J., Vangel, M., Congleton, C., Yerramsetti, S. M., Gard, T., & Lazar, S. W. (2011). Mindfulness practice leads to increases in regional brain gray matter density. Psychiatry Research, 191(1), 36–43. 10.1016/j.pscychresns.2010.08.006.

Hölzel, B. K., Ott, U., Gard, T., Hempel, H., Weygandt, M., Morgen, K., & Vaitl, D. (2008). Investigation of mindfulness meditation practitioners with voxel-based morphometry. Social Cognitive and Affective Neuroscience, 3(1), 55–61. 10.1093/scan/nsm038.

JASP Team (2023). JASP (Version 0.17.2) [Computer software]. https://jasp-stats.org.

Kane, M. J., Brown, L. E., Little, J. C., Silvia, P. J., Myin-Germeys, I., & Kwapil, T. R. (2007). For whom the mind wanders, and when: An experience-sampling study of working memory and executive control in daily life. Psychological Science, 18, 614– 621. 10.1111/j.1467-9280.2007.01948.x.

Kane, M. J., & McVay, J. C. (2012). What mind wandering reveals about executive-control abilities and failures. Current Directions in Psychological Science, 21(5), 348–354. 10.1177/0963721412454875.

Killingsworth, M. A., & Gilbert, D. T. (2010). A wandering mind is an unhappy mind. Science, 330(6006), 932. 10.1126/science.1192439.

Kucyi, A., Kam, J. W. W., Andrews-Hanna, J. R., Christoff, K., & Whitfield-Gabrieli, S. (2023). Recent advances in the neuroscience of spontaneous and off-task thought: Implications for mental health. Nature Mental Health, 1(11), 827–840. 10.1038/s44220-023-00133-w.

Liu, J., & Calhoun, V. (2007). Parallel independent component analysis for multimodal analysis: Application to fMRI and EEG data. In 2007 4th IEEE International Symposium on Biomedical Imaging: From Nano to Macro (pp. 1028-1031). IEEE. 10.1109/isbi.2007.357030.

Lu, H., Song, Y., Xu, M., Wang, X., Li, X., & Liu, J. (2014). The brain structure correlates of individual differences in trait mindfulness: A voxel-based morphometry study. Neuroscience, 272, 21–28. 10.1016/j.neuroscience.2014.04.051.

Luders, E., Toga, A. W., Lepore, N., & Gaser, C. (2009). The underlying anatomical correlates of long-term meditation: Larger hippocampal and frontal volumes of gray matter. NeuroImage, 45(3), 672–678. 10.1016/j.neuroimage.2008.12.061.

Mas-Cuesta, L., Baltruschat, S., Cándido, A., Verdejo-Lucas, C., Catena-Verdejo, E., & Catena, A. (2024). Brain changes following mindfulness: Reduced caudate volume is associated with decreased positive urgency. Behavioural Brain Research, 461, 114859. 10.1016/j.bbr.2024.114859.

Mason, M. F., Norton, M. I., Van Horn, J. D., Wegner, D. M., Grafton, S. T., & Macrae, C. N. (2007). Wandering minds: The default network and stimulus-independent thought. Science, 315(5810), 393–395. 10.1126/science.1131295.

McNab, F., & Klingberg, T. (2008). Prefrontal cortex and basal ganglia control access to working memory. Nature Neuroscience, 11(1), 103–107. 10.1038/nn2024.

McVay, J. C., & Kane, M. J. (2010). Does mind wandering reflect executive function or executive failure? Comment on Smallwood and Schooler (2006) and Watkins (2008). Psychological Bulletin, 136(2), 188–197. 10.1037/a0018298.

Melis, M., Schroyen, G., Pollefeyt, J., Raes, F., Smeets, A., Sunaert, S., Deprez, S., & Van der Gucht, K. (2022) The impact of mindfulness-based interventions on brain functional connectivity: A systematic review. Mindfulness, 13(8), 1857–1875. 10.1007/s12671-022-01919-2.

Mendes, N., Oligschläger, S., Lauckner, M. E., Golchert, J., Huntenburg, J. M., Falkiewicz, M., Ellamil, M., Krause, S., Baczkowski, B. M., Cozatl, R., Osoianu, A., Kumral, D., Pool, J., Golz, L., Dreyer, M., Haueis, P., Jost, R., Kramarenko, Y., Engen, H., … Margulies, D. S. (2019). A functional connectome phenotyping dataset including cognitive state and personality measures. Scientific Data, 6, 180307. 10.1038/sdata.2018.307.

Messina, I., Grecucci, A., & Viviani, R. (2023). Neurobiological models of emotion regulation: A meta-analysis of neuroimaging studies of acceptance as an emotion regulation strategy. Social Cognitive and Affective Neuroscience, 16(3), 257–267. 10.1093/scan/nsab007.

Modinos, G., Ormel, J., & Aleman, A. (2010). Individual differences in dispositional mindfulness and brain activity involved in reappraisal of emotion. Social Cognitive and Affective Neuroscience, 5(4), 369–377. 10.1093/scan/nsq006.

Monachesi, B., Grecucci, A., Ghomroud, P. A., & Messina, I. (2023). Comparing reappraisal and acceptance strategies to understand the neural architecture of emotion regulation: A meta-analytic approach. Frontiers in Psychology, 14, 1187092. 10.3389/fpsyg.2023.1187092.

Mooneyham, B. W., & Schooler, J. W. (2013). The costs and benefits of mind-wandering: A review. Revue Canadienne de Psychologie Expérimentale, 67(1), 11–18. 10.1037/a0031569.

Mrazek, M. D., Franklin, M. S., Phillips, D. T., Baird, B., & Schooler, J. W. (2013a). Mindfulness training improves working memory capacity and GRE performance while reducing mind wandering. Psychological Science, 24(5), 776–781. 10.1177/0956797612459659.

Mrazek, M. D., Phillips, D. T., Franklin, M. S., Broadway, J. M., & Schooler, J. W. (2013b). Young and restless: Validation of the Mind-Wandering Questionnaire (MWQ) reveals disruptive impact of mind-wandering for youth. Frontiers in Psychology, 4, 560. 10.3389/fpsyg.2013.00560.

Murakami, H., Nakao, T., Matsunaga, M., Kasuya, Y., Shinoda, J., Yamada, J., & Ohira, H. (2012). The structure of mindful brain. PLoS ONE, 7(9), Article e46377. 10.1371/journal.pone.0046377.

Ottaviani, C., & Couyoumdjian, A. (2013). Pros and cons of a wandering mind: A prospective study. Frontiers in Psychology, 4, Article 524. 10.3389/fpsyg.2013.00524.

Packard, M. G., & Knowlton, B. J. (2002). Learning and memory functions of the basal ganglia. Annual Review of Neuroscience, 25, 563–593. 10.1146/annurev.neuro.25.112701.142937

Parkinson, T. D., Kornelsen, J., & Smith, S. D. (2019). Trait mindfulness and functional connectivity in cognitive and attentional resting state networks. Frontiers in Human Neuroscience, 13, 112. 10.3389/fnhum.2019.00112.

Penny, W. D., Friston, K. J., Ashburner, J. T., Kiebel, S. J., & Nichols, T. E. (Eds.). (2011). Statistical parametric mapping: The analysis of functional brain images. Elsevier.

Posner, M. I., Tang, Y. Y., & Lynch, G. (2014). Mechanisms of white matter change induced by meditation training. Frontiers in Psychology, 5, 1220. 10.3389/fpsyg.2014.01220.

R Core Team. (2021). R: A language and environment for statistical computing. R Foundation for Statistical Computing, Vienna, Austria. https://www.R-project.org.

Seli, P., Carriere, J. S., & Smilek, D. (2014). Not all mind wandering is created equal: Dissociating deliberate from spontaneous mind wandering. Psychological Research, 79, 750–758. 10.1007/s00426-014-0617-x.

Seli, P., Risko, E. F., & Smilek, D. (2016a). Assessing the associations among trait and state levels of deliberate and spontaneous mind wandering. Consciousness and Cognition: An International Journal, 41, 50–56. 10.1016/j.concog.2016.02.002.

Seli, P., Risko, E. F., Smilek, D., & Schacter, D. L. (2016b). Mind-wandering with and without intention. Trends in Cognitive Sciences, 20(8), 605–617. 10.1016/j.tics.2016.05.010.

Smallwood, J., Davies, J. B., Heim, D., Finnigan, F., Sudberry, M. V., O’Connor, R., & Obonsawin, M. C. (2004). Subjective experience and the attentional lapse: Task engagement and disengagement during sustained attention. Consciousness and Cognition, 13, 657–690. 10.1016/j.concog.2004.06.003.

Smallwood, J., O’Connor, R. C., Sudbery, M. V., & Obonsawin, M. (2007). Mind-wandering and dysphoria. Cognition and Emotion, 21(4), 816–842. 10.1080/02699930600911531.

Smallwood, J., & Schooler, J. W. (2006). The restless mind. Psychological Bulletin, 132(6), 946. 10.1037/0033-2909.132.6.946.

Smallwood, J., & Schooler, J. W. (2015). The science of mind wandering: Empirically navigating the stream of consciousness. Annual Review of Psychology, 66, 487–518. 10.1146/annurev-psych-010814-015331.

Sorella, S., Crescentini, C., Matiz, A., Chang, M., & Grecucci, A. (2025). Resting-state BOLD temporal variability of the default mode network predicts spontaneous mind wandering, which is negatively associated with mindfulness skills. Frontiers in Human Neuroscience, 19, Article 1515902. 10.3389/fnhum.2025.1515902.

Tang, R., Friston, K. J., & Tang, Y. Y. (2020). Brief mindfulness meditation induces gray matter changes in a brain hub. Neural Plasticity, 2020, 8830005. 10.1155/2020/8830005.

Tang, Y. Y., Hölzel, B. K., & Posner, M. I. (2015). The neuroscience of mindfulness meditation. Nature Reviews Neuroscience, 16(4), 213–225. 10.1038/nrn3916.

Tang, Y.-Y., Lu, Q., Fan, M., Yang, Y., & Posner, M. I. (2012). Mechanisms of white matter changes induced by meditation. Proceedings of the National Academy of Sciences, 109(26), 10570–10574. 10.1073/pnas.1207817109.

Tang, Y. Y., Lu, Q., Geng, X., Stein, E. A., Yang, Y., & Posner, M. I. (2010). Short-term meditation induces white matter changes in the anterior cingulate. Proceedings of the National Academy of Sciences, 107(35), 15649–15652. 10.1073/pnas.1011043107.

Taren, A. A., Creswell, J. D., & Gianaros, P. J. (2013). Dispositional mindfulness co-varies with smaller amygdala and caudate volumes in community adults. PloS ONE, 8(5), e64574. 10.1371/journal.pone.0064574.

Taylor, V. A., Grant, J., Daneault, V., Scavone, G., Breton, E., Roffe-Vidal, S., Courtemanche, J., Lavarenne, A. S., & Beauregard, M. (2011). Impact of mindfulness on the neural responses to emotional pictures in experienced and beginner meditators. NeuroImage, 57(4), 1524–1533. 10.1016/j.neuroimage.2011.06.001.

The MathWorks Inc. (2022). MATLAB version: 2018b, Natick, Massachusetts: The MathWorks Inc. https://www.mathworks.com.

Vestergaard-Poulsen, P., van Beek, M., Skewes, J., Bjarkam, C. R., Stubberup, M., Bertelsen, J., & Roepstorff, A. (2009). Long-term meditation is associated with increased gray matter density in the brain stem. Neuroreport, 20(2), 170–174. 10.1097/WNR.0b013e328320012a.

Wang, X., Xu, M., Song, Y., Li, X., Zhen, Z., Yang, Z., & Liu, J. (2014). The network property of the thalamus in the default mode network is correlated with trait mindfulness. Neuroscience, 278, 291–301. 10.1016/j.neuroscience.2014.08.006.

Ward, L. M. (2013). The thalamus: Gateway to the mind. Wiley Interdisciplinary Reviews: Cognitive Science, 4(6), 609–622. 10.1002/wcs.1256.

Wax, M., & Kailath, T. (1985). Detection of signals by information theoretic criteria. IEEE Transactions on Acoustics, Speech, and Signal Processing, 33(2), 387–392. 10.1109/TASSP.1985.1164557.

Yuan, S., Wu, H., Wu, Y., Xu, H., Yu, J., Zhong, Y., Zhang, N., Li, J., Xu, Q., & Wang, C. (2022). Neural effects of cognitive behavioral therapy in psychiatric disorders: A systematic review and activation likelihood estimation meta-analysis. Frontiers in Psychology, 13, 853804. 10.3389/fpsyg.2022.853804.

Zhang, J., Raya, J., Morfini, F., Urban, Z., Pagliaccio, D., Yendiki, A., Auerbach, R. P., Bauer, C. C. C., & Whitfield-Gabrieli, S. (2023). Reducing default mode network connectivity with mindfulness-based fMRI neurofeedback: A pilot study among adolescents with affective disorder history. Molecular Psychiatry, 28, 2540–2548. 10.1038/s41380-023-02032-z.

Zhuang, K., Bi, M., Li, Y., Xia, Y., Guo, X., Chen, Q., Du, X., Wang, K., Wei, D., Yin, H., & Qiu, J. (2017). A distinction between two instruments measuring dispositional mindfulness and the correlations between those measurements and the neuroanatomical structure. Scientific Reports, 7, Article 6252. 10.1038/s41598-017-06599-w.

